# Rapid and Highly Efficient Morphogenic Gene-Mediated Hexaploid Wheat Transformation

**DOI:** 10.1101/2023.01.25.525570

**Authors:** Kari Johnson, Uyen Cao Chu, Geny Anthony, Emily Wu, Ping Che, Todd J. Jones

## Abstract

The successful employment of morphogenic regulator genes, *Zm-Baby Boom* (*ZmBbm*) and *Zm-Wuschel2* (*ZmWus2*), for *Agrobacterium*-mediated transformation of maize (*Zea mays* L.) and sorghum (*Sorghum bicolor* L.) has been reported to improve transformation by inducing rapid somatic embryo formation. Here, we report two morphogenic gene-mediated wheat transformation methods, either with or without morphogenic and marker gene excision. These methods yield independent-transformation efficiency up to 58% and 75%, respectively. In both cases, the tissue culture duration for generating transgenic plants was significantly reduced from 80 to nearly 50 days. In addition, the transformation process was significantly simplified to make the procedure less labor-intensive, higher-throughput and more cost-effective, by eliminating the requirement for embryonic axis excision, bypassing the necessity for prolonged dual-selection steps for callus formation, and obviating the prerequisite of cytokinin for shoot regeneration. Furthermore, we have demonstrated the flexibility of the methods and generated high-quality transgenic events across multiple genotypes using herbicide (phosphinothricin, ethametsulfuron)- and antibiotic (G418)-based selections.

## INTRODUCTION

Wheat (*Triticum aestivum* L.) is a staple crop around the world but engineering its genome has historically been challenging due to its polyploid nature (Shewry, 2009; Borrill et al., 2019; Li et al., 2021). Robust transformation technologies coupled with efficient gene editing is essential for accelerated breeding and the development of superior varieties. However, conventional wheat transformation (**Figure3A**) has been known for its genotype dependency (Harwood, 2012) and its costly, time-consuming, and labor-intensive tissue culture procedure, such as the excessive explant manipulation to remove the embryonic axis and extensive dual-selection steps for transgenic callus formation (Ishida et al., 2015; Hayta et al., 2019) (**Figure3A**). Therefore, developing a simple, reproducible, and more efficient wheat transformation system that overcomes genotype-dependent barriers is critical for wheat genetic improvement through gene integration and CRISPR/Cas-mediated genome-modification.

Recently, significant progress has been made toward more efficient wheat transformation with reduced genotype dependency and tissue culture cycle time by application of growth-regulating and regeneration related genes, such as *TaWox5* (Wang et al., 2022) and the chimeric fusion *Grf-Gif* (Debernardi et al., 2020). In this study, we have taken advantage of the *Zm-Baby Boom* (*ZmBbm*) and *Zm-Wuschel2* (*ZmWus2*) mediated transformation technologies developed in maize (*Zea mays* L.) (Lowe et al., 2016) and sorghum (*Sorghum bicolor* L.) (Che et al., 2022). We extended those technologies to wheat and established two wheat transformation methods (**Figure 3B**) for generating high-quality events across multiple genotypes, either with or without morphogenic and marker gene excision (hereby referred to as the ‘QuickWheat’ transformation system). The non-excision transformation method generated transgenic events with integrated morphogenic genes and selectable marker genes, a method suitable for genome editing applications. The excision transformation method generated transgenic events without morphogenic and selectable marker genes through *moCRE/loxP*-mediated gene cassette excision, an advanced, well-suited approach for trait gene function characterization. In both cases, due to the rapid somatic embryo formation induced by *ZmWus2* and/or *ZmBbm*, several procedures considered essential for conventional wheat transformation were eliminated. For example, the prerequisite of cytokinins in regeneration medium for shoot induction and elongation was removed. The requirements of embryonic axis excision after co-cultivation and the prolonged dual-selection steps for callus formation were eliminated to reduce the workload as well. The total time from inoculation of immature embryos to transplantation of a fully developed transgenic plant in the greenhouse was reduced to nearly 50 days (**Figure 3B**) compared to around 80 days (**Figure 3A**) for conventional wheat transformation reported previously (Ishida et al., 2015; Hayta et al., 2019). Furthermore, we have demonstrated the flexibility of the QuickWheat system and generated high-quality transgenic events across multiple genotypes using different antibiotic- and herbicide-based selections.

## MATERIALS AND EQUIPMENT

1. Laminar flow hood, with sterilizing unit and beads
2. Personal protective equipment (safety glasses, lab coat)
3. *Agrobacterium* LBA4404 Thy- containing construct
4. Medium #25 (**Table 1**)
5. Medium #50 (**Table 1**)
6. Glass petri dish 100×250 mm
7. Wheat variety SBC0456D, or other spikes (**Figure 1A**)
8. Corning stir plate and magnetic stir bar
9. Bleach (8.25% NaOCl)
10. Tween-20
11. Sterile water
12. Cylindrical beaker
13. Mesh strainer
14. Petri dish (150×15 mm)
15. Embryo isolation tool (wax carving tool or other similar with a flat face) (**Supplementary Figure 1B**)
16. Embryo orienting tool (Hu-Friedy TNPF18A or other similar)
17. VWR Round/Tapered microspatula tool or other similar
18. Infection tube, Fisher Scientific, sterile 2.0 mL conical microcentrifuge screw cap tube
19. 15-50 mL plastic tubes
20. Pipettes and pipette tips (10-5000 μL)
21. Sterile loops
22. Culture boxes
23. Acetosyringone 400 mM (Sigma-Aldrich)
24. Medium #100 (**Table 2**)
25. Medium #200 (**Table 2**)
26. Medium #300 (**Table 2**)
27. Medium #400 (**Table 2**)
28. Stereomicroscope
29. VWR mini-incubator
30. Eppendorf miniSpin plus centrifuge (**Supplementary Figure 1A**)
31. Thermo Scientific Genesys30 Visible Spectrophotometer
32. Intellus Percival environmental controller at 21°C
33. Dark culture room at 28°C
34. Bright light culture room at 26°C, 16hr photoperiod, Valoya C65 NS12 LED light set (BX Series, Valoya, Finland) set to 100 μmol/m^2^/s

**TABLE 1.**
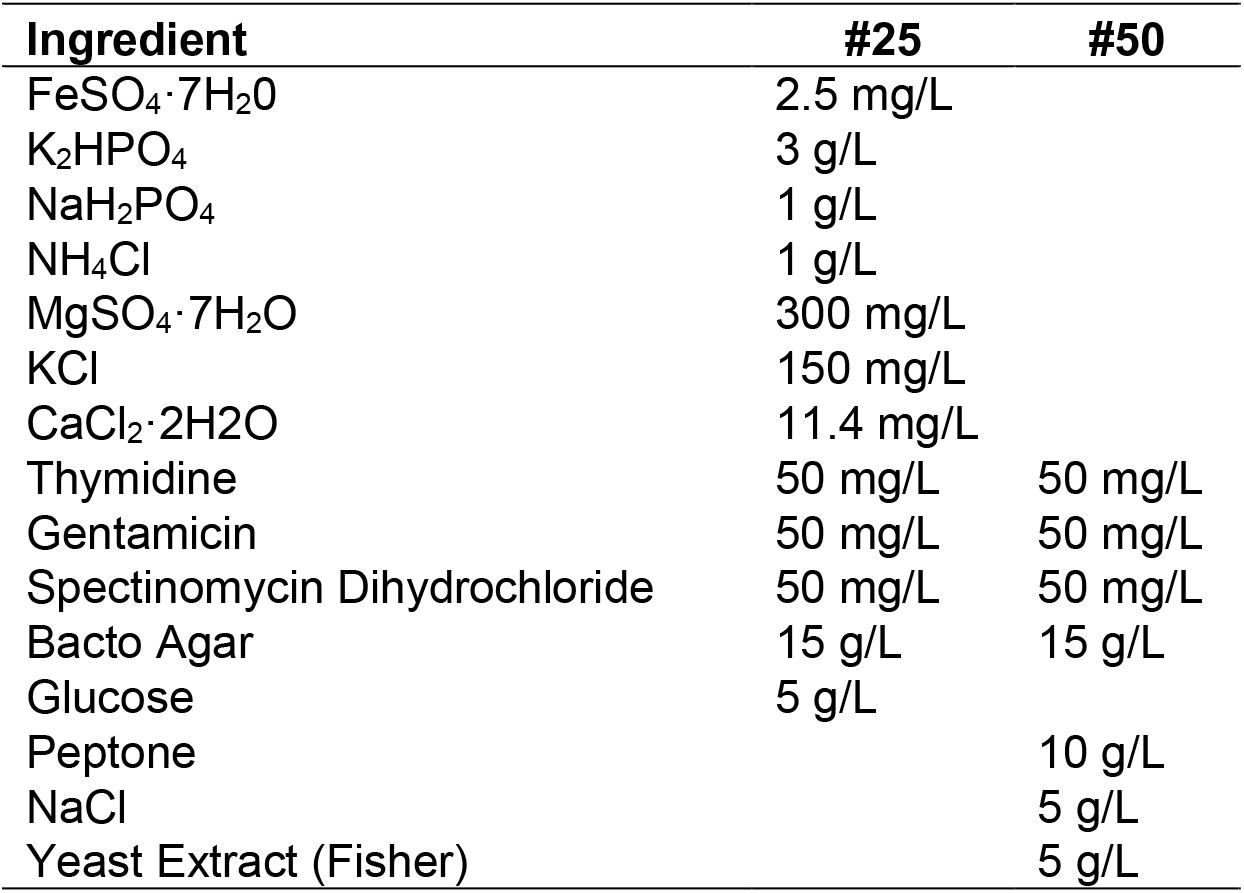
Medium recipes for master and working plates

**TABLE 2.**
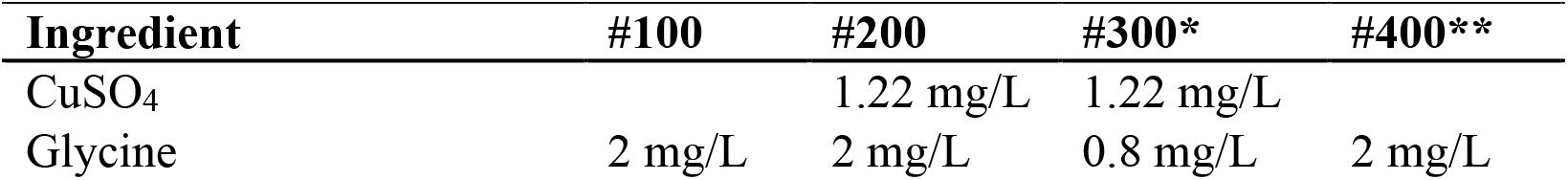
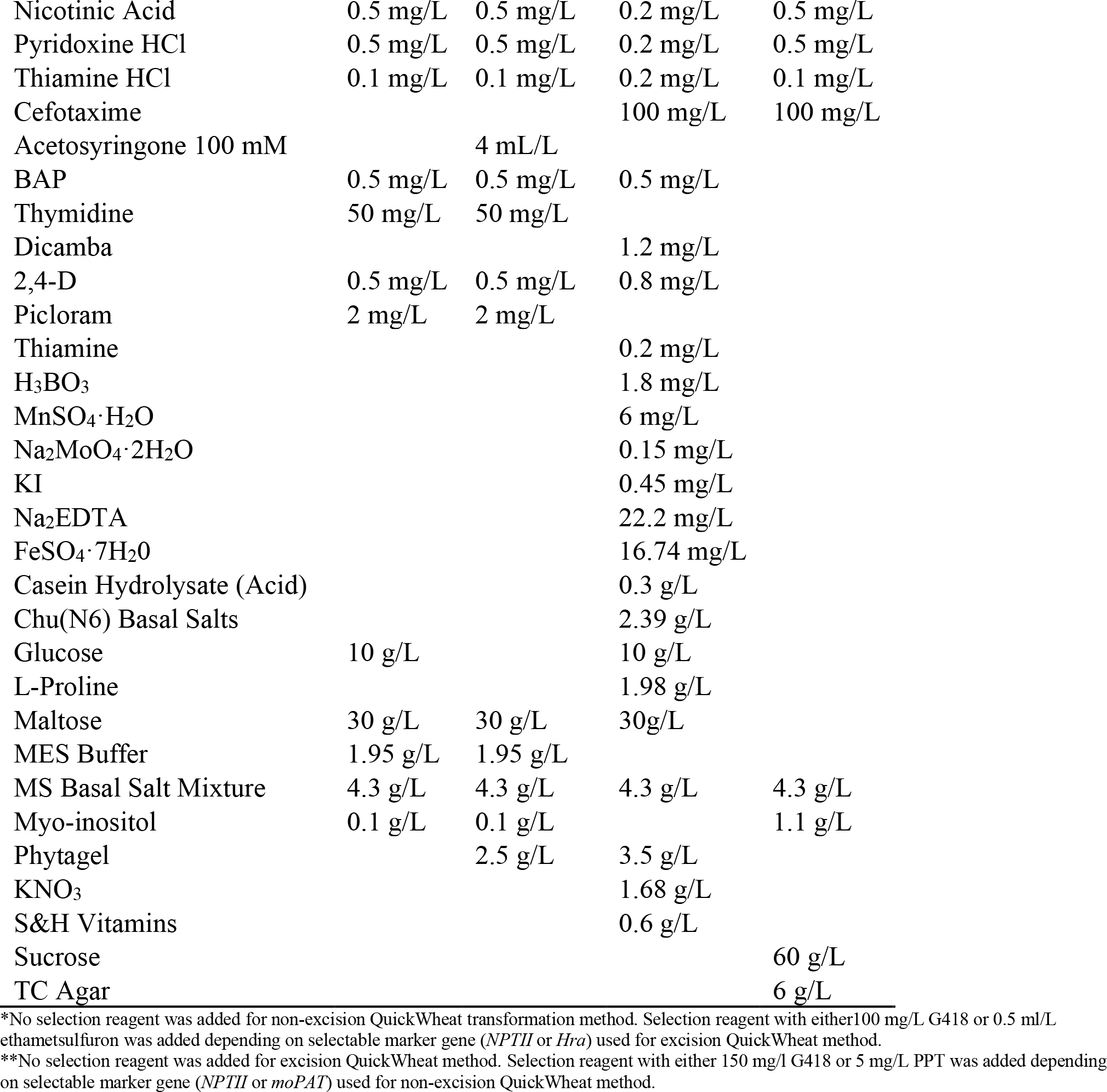
Medium recipes for transformation

**FIGURE 1.**
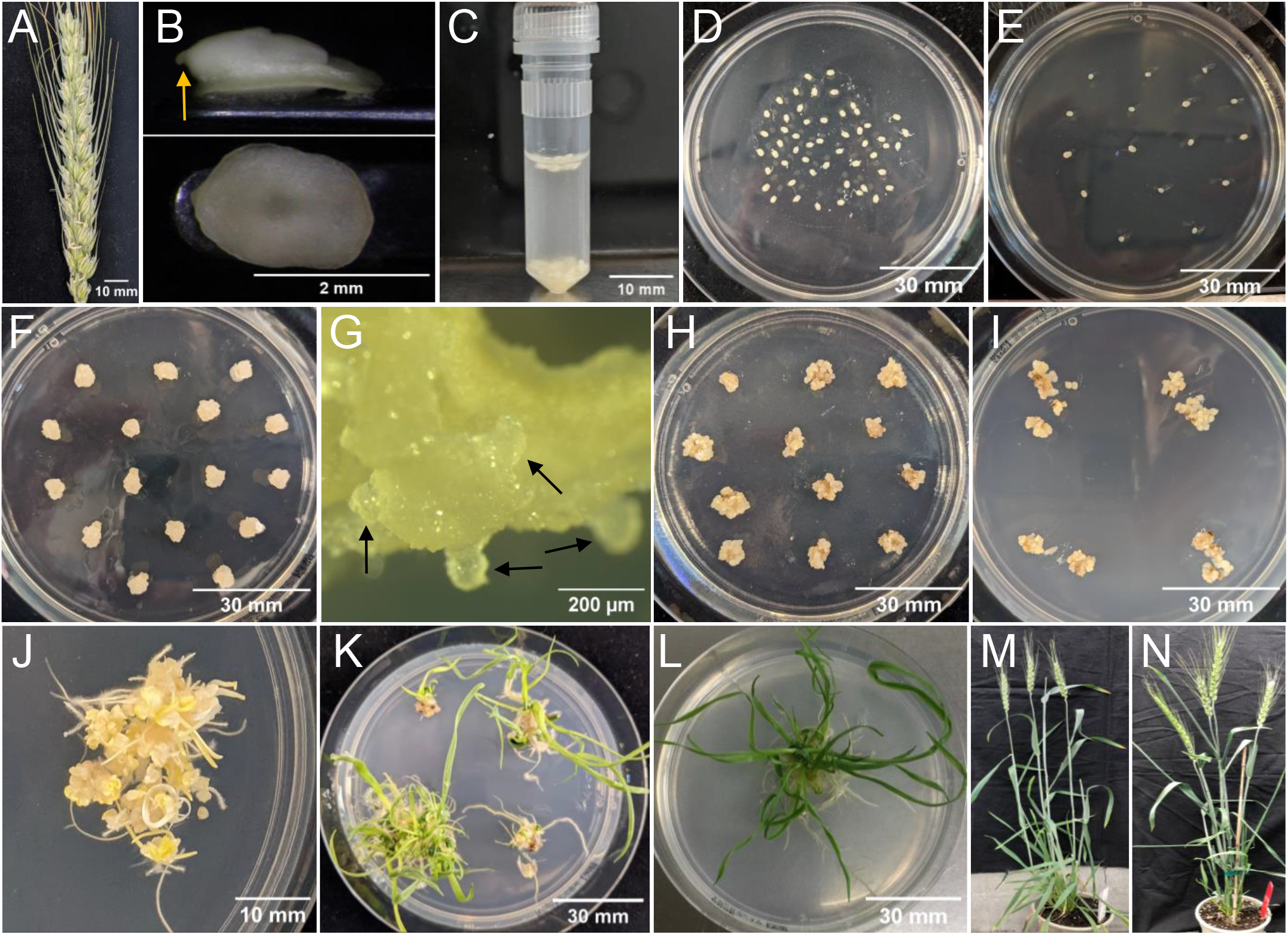
Tissue culture response using morphogenic genes in spring wheat SBC0456D. (**A**) Selected spike at 14 days post-anthesis. (**B**) Isolated immature embryo ranging from 1.8-2.1 mm in length. The top photo shows the embryo in profile, with an arrow pointing to the radicle. The bottom photo shows the scutella face of the embryo. (**C**) Approximately 50 immature embryos in an infection tube prior to centrifugation. (**D**) Immature embryos with scutellum side up, on co-cultivation medium immediately following infection. (**E**) Immature embryos with scutellum side up on resting medium, one day after co-cultivation. (**F**) Callus induction of immature embryos at the end of resting stage. (**G**) Close look of somatic embryo formation (few examples indicated by the arrows) at the end of resting stage. (**H**) Proliferation of transformed tissue at the end of callus induction with selection before heat shock treatment. (**I**) Embryogenic callus tissue from four immature embryos were broken into several pieces and putted onto regeneration and elongation medium either with or without selection based on QuickWheat transformation methods. (**J**) Shoot and root initiation on regeneration medium after 2 weeks in the dark. (**K**) Regeneration and elongation of shoots and roots after 14 days under LED light before plantlets were separated. (**L**) Separated and fully developed individual T0 plant ready to be sent to controlled environments. (**M**) Morphology of wild-type plant germinated from seed. (**N**) Normal and fertile T0 event carrying two copies of *ZmWUS2* and *ZmBbm*. (**A-G**) Photos representing the common tissue culture stages for both non-excision and excision wheat transformation methods. (**H**) Photo representing the unique tissue culture stage for excision wheat transformation method. (**I-L**) Photos representing the relatively similar tissue culture stages for both non-excision and excision wheat transformation methods.

## METHODS

Two QuickWheat transformation methods, non-excision and excision, were developed as illustrated in **Figure 3B**. The non-excision transformation method generated transgenic events with integrated morphogenic genes and a selectable marker gene. The excision transformation method generated morphogenic gene free and marker gene free transgenic events through *moCRE/loxP*-mediated gene cassette excision. The two transformation methods shared some identical steps, such as *Agrobacterium* inoculation, co-cultivation and resting, during the early stage of the transformation process. However, unique procedures were applied after the resting step based on the transformation method. The transformation procedures for each of the transformation methods are described in detail below and illustrated in **Figure 3B**.

### Growing Donor Plants to Produce Immature Embryos for Transformation

1. Wheat genotypes, SBC0456D, Fielder, and Chinese Spring, grown in a greenhouse and growth chamber were used in this study. Plant one wheat seed into an ellepot™ plug (Ellepot A/S, Esbjerg, Denmark). Set two ellepot plugs into each pot of substrate consisting of peat, bark, perlite, wetting agent, lime, starter and silicone. Place pots in the greenhouse for two weeks with a photoperiod of 16 hours and daytime temperatures ranging from 27°C to 21°C. Vapor pressure deficit ranges (VPD) from 10-14millibars (mb) and daily light integral (DLI) 28-38 moles/m^2^/day.
2. Move pots to a growth chamber and transplant plants into new substrate mix (peat, perlite, wetting agent, lime, starter and silicone). Set growth chamber to 22°C during the day and 16°C over night, with 55% relative humidity, 350-400 μmol/m^2^/s lighting, and 16-hour photoperiod.
3. Bag spikes at the start of flowering. Fourteen days later, or when embryos are 1.8-2.1 mm long, harvest entire spike.

### *Agrobacterium* Preparation

1. The constructs used in this experiment are illustrated in **Figure 2**.
2. An *Agrobacterium* auxotrophic strain of LBA4404 Thy-containing a pPHP71539 (pVir) ternary vector was frozen as a glycerol stock (Anand et al., 2018; Che et al., 2018).
3. Streak *Agrobacterium* from the glycerol stock onto a master plate medium #25 (**Table 1**). Incubate 3-4 days at 28°C in the dark. The cultured *Agrobacterium* on the master plate can be stored in 4°C refrigerator and used for up to one month.
4. About 16-20 hours before transformation initiation, streak the cultured *Agrobacterium* from the master plate to the working plate medium #50 (**Table 1**). Incubate at 28°C in the dark.

**FIGURE 2.**
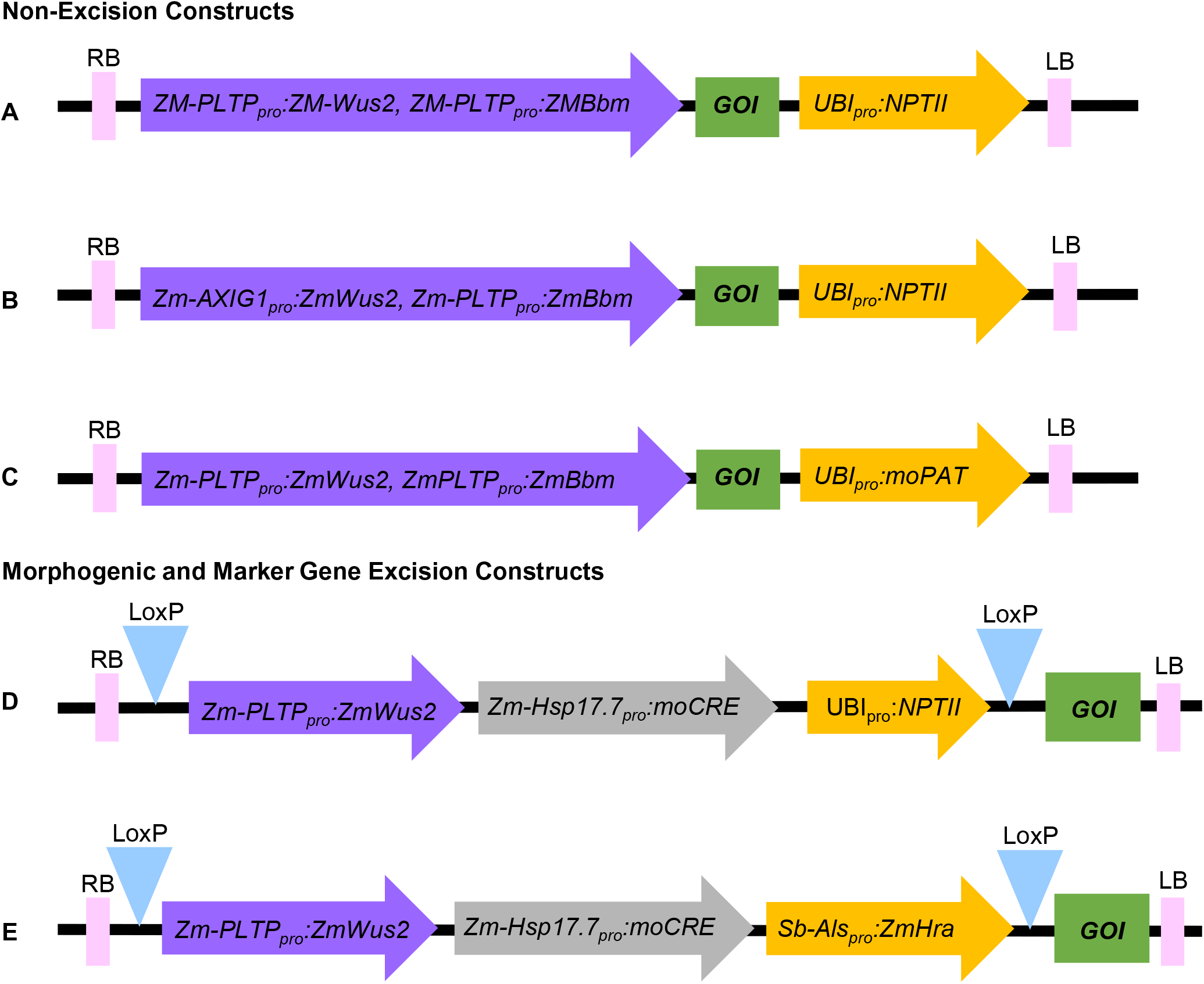
Schematic representation of the molecular components of constructs used in this study. (**A**) and (**B**) Non-excision constructs using *NPTII* as selectable marker. (**C**) Non-excision construct using *moPAT* as selectable marker. (**D**) Morphogenic and marker gene excision construct using *NPTII* as selectable marker. (**E**) Morphogenic and marker gene excision construct using *Hra* as selectable marker. RB: Right border, LB: Left border, GOI: Gene of interest.

**FIGURE 3.**
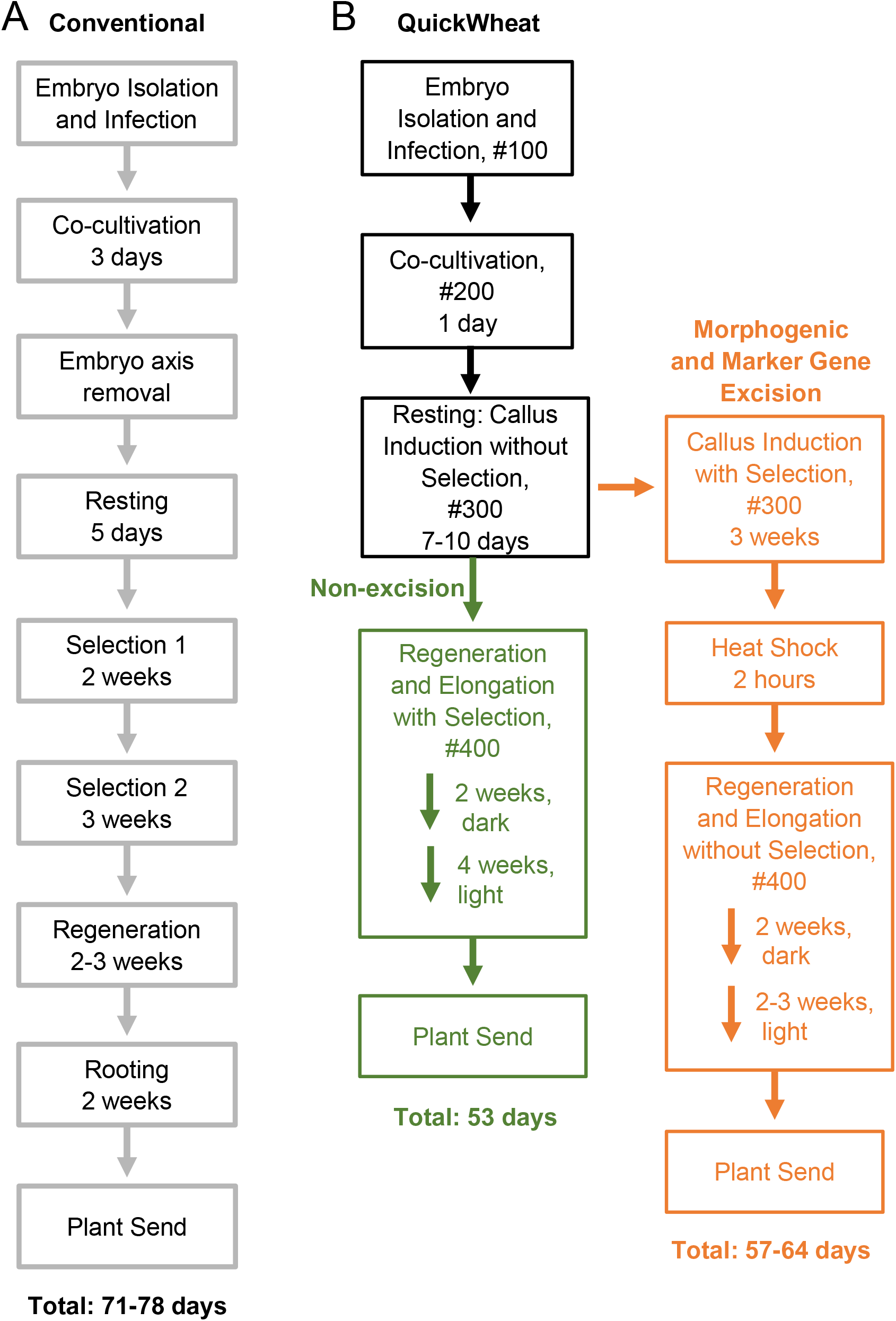
Flow chart comparing conventional wheat transformation and QuickWheat transformation procedures. (**A**) Conventional transformation procedure (**B**) QuickWheat transformation procedure either with or without morphogenic and marker gene excision.

### Seed Sterilization and Infection Preparation

1. Collect spikes containing immature seeds (with 1.8-2.1 mm embryos) (**Figure 1A and B**). Remove seeds from spikes and place in glass petri dish.
2. Sterilize seeds in glass petri dish with 20% bleach solution and two drops of Tween-20 for 15 minutes. Use a magnetic stir plate and bar to stir on level 3-6.
3. Remove bleach and Tween-20 using a mesh strainer. Rinse seeds with sterile water.
4. Let the seeds sit for five minutes in sterile water in the glass petri dish.
5. Remove water using the mesh strainer and place the sterile seeds in a large petri dish (150×15 mm).
6. In a 50 mL tube prepare initiation solution by adding 1 μL of freshly thawed 400 mM acetosyringone (AS) for every 1 mL of #100 liquid medium (**Table 2**). Shake to mix.
7. Distribute 2 mL of initiation solution into as many infection tubes as needed.

### Immature Embryo Pre-treatment, Infection, Co-Cultivation and Callus Induction

1. Immature embryo isolation, pre-treatment, and infection: Using an embryo isolation tool (**Supplementary Figure 1B**), isolate 1.8-2.1 mm embryos from the sterilized wheat seeds (**Figure 1B**) and put them in the infection tube containing initiation solution. Each infection tube should contain 30-60 embryos when isolating embryos is complete (**Figure 1C**).
2. Centrifuge the infection tube at 6,000 RPM using Eppendorf miniSpin plus centrifuge (**Supplementary Figure 1A**) for five minutes at room temperature.
3. In a 15 mL tube prepare *Agrobacterium* solution using the initiation solution as described above. Suspend several loops of the *Agrobacterium* culture from the working plate into the initiation solution and make a homogenous suspension. Adjust the O.D. of the *Agrobacterium* suspension to 1.0 at 600 nm.
4. Once centrifugation of the infection tube is complete use a pipette to remove only the initiation solution before adding 1 mL of *Agrobacterium* suspension (from Step 3) to the embryos.
5. Let the embryos sit in the *Agrobacterium* suspension for five minutes.
6. Rotate the infection tube to dislodge the embryos. Pour out the *Agrobacterium* suspension and embryos quickly onto co-cultivation medium #200 (**Table 2**). Use a microspatula to remove any embryos from the infection tube if they stick to the sides or lid.
7. Co-Cultivation: Using a microscope to arrange the embryos scutellum side up. Make sure the embryos are immersed in *Agrobacterium* suspension, touching the co-cultivation medium, and evenly spaced out. Incubate in the dark at 21°C for one day (**Figure 1D**).
8. Resting (callus induction without selection): After one day on co-cultivation, the embryos remain fragile. Carefully transfer 10-20 embryos scutellum side up to medium #300 (**Table 2**) for callus induction. Push the radicle of the embryo (**Figure 1B**) into the medium to ensure they do not begin to germinate (**Figure 1E**). Incubate in a dark culture room for seven to ten days. Throughout this time, the embryos approximately quadruple in size.

### Shoot and Root Regeneration and Elongation of Transgenic Plants

Depending on the transformation method of the QuickWheat system (excision or non-excision), the tissue culture process for selection, heat-shock treatment, and regeneration and elongation was conducted differently after the resting step. The differences for each transformation method are described below.

#### For non-excision transformation method

1. Regeneration of shoots and roots with selection: After resting for ten days (**Figure 1F**), move tissue to regeneration medium #400 (**Table 2**) with either 150 mg/L G418 or 5 mg/L PPT added to the medium depending on selectable marker gene (*NPTII* or *moPAT*). Break up the tissue into 3-5 pieces and spread a bit on the medium to allow room for growth (**Figure 1I**). Transfer up to four embryos per plate. Incubate in dark culture room for two weeks. Following two weeks in the dark room, 50-100% of embryos begin to form shoots and roots (**Figure 1J**).
2. Elongation of shoots and roots: After two weeks in a dark, move tissue to a bright light culture room. After one to two weeks in the light, regenerated shoot and root growth are observed. Shoots elongate and turn green with significant development. Primary roots are a few centimeters long and begin to form secondary roots (**Figure 1K**). Separate and transfer each plantlet onto a new plate with medium #400 (**Table 2**) with G418 or PPT for elongation with *NPTII* or *moPAT* as selectable marker (depending on selectable marker gene). Allow plants to continue growing, approximately two weeks, until sending T0 plants to the greenhouse (**Figure 1L and N**).

#### For excision transformation method

1. Callus induction with selection: After resting for seven days (**Figure 1F**), move tissue to medium #300 (**Table 2**) with either 100 mg/L G418 or 0.5 ml/L ethametsulfuron depending on selectable marker gene (*NPTII* or *Hra*) for three weeks in dark culture room (**Figure 1H**).
2. Heat shock treatment: Place culture boxes with tissue in Percival chamber at 45°C with 70% relative humidity for 2 hours.
3. Regeneration of shoots and roots: After removal from heat shock treatment allow the plates to cool down to room temperature (roughly two hours). Transfer tissue to medium #400 (**Table 2**) for regeneration of shoots and roots. Break up the tissue into 3-5 pieces and spread a bit on medium to allow room for growth (**Figure 1I**). Transfer up to four embryos per plate of medium #400 (**Table 2**). Incubate in a dark culture room for two weeks. Following two weeks in the dark room, 50-100% of embryos begin to form shoots and roots (**Figure 1J**).
4. Elongation of shoots and roots: After two weeks in the dark, move tissue to a bright light culture room. After one to two weeks in the light, regenerated shoot and root growth are observed. Shoots elongate and turn green with significant development. Primary roots are a few centimeters long and begin to form secondary roots (**Figure 1K**). Separate and transfer each plantlet onto a new plate with medium #400 (**Table 2**). Allow plants to continue growing, approximately one week, until sending T0 plants to the greenhouse (**Figure 1L**).

### T0 Transplant in Controlled Environments (CE)

1. Transplant each T0 from its tissue culture plate into an ellepot plug and set into substrate consisting of peat, perlite, wetting agent, lime, starter fertilizer and silicone. Place plugs in flats in a greenhouse with a photoperiod of 16 hours and daytime temperatures ranging from 21°C to 27°C. VPD ranges from 10-14mb and DLI 28-38 moles/day. Leaf sample plants seven days after being received by CE.
2. After two to three weeks in the greenhouse move pots to a growth chamber or new greenhouse and transplant plants into new substrate mix (peat, bark, perlite, wetting agent, lime, starter fertilizer and silicone. Set greenhouse at 22°C during the day and 15-17°C over night, DLI 25 moles/day with 16-hour photoperiod. T0 survival after sending being received by the CE is greater than 99%.

### Event Quality and Dependency Analysis

Stable T-DNA integration and event quality were determined through a series of quantitative PCR (qPCR) analyses using genomic DNA extracted from the putative T0 transgenic events based on the method previously described (Wu et al., 2014; Lowe et al., 2016; Hoerster et al., 2020). In brief, as shown in **Supplementary Figure 2**, nine qPCR assays were developed to determine T-DNA integrity or intactness. Three of these assays (PSA2, PSB1 and NPTII) were used for copy number determination and six assays (ZmWUS2, PSW1, ZmBBM, PSO1, PSN1, PSN2) for presence/absence of the T-DNA elements. Outside the border integration sites, PCR backbone-specific assays were developed to check for any border read-through (Wu et al., 2014; Lowe et al., 2016; Hoerster et al., 2020). The presence or absence of *Agrobacterium* vector backbone integration of the binary vector was detected based on screening for sequences from four regions outside of the T-DNA integration sites of the vector, such as LB, REP A, REP B and REP C. Based on those analysis, the transgenic plant carrying a single copy of the intact T-DNA integrations and without vector backbone was defined as a ‘quality event (QE)’ for non-excision transformation, and a transgenic plant carrying a single copy of the intact T-DNA integrations without vector backbone and ZmWus2/moCRE/selectable-marker cassette was defined as a ‘quality event (QE)’ for excision transformation. All other events were defined as ‘non-quality event (non-QE)’

## RESULTS AND DISCUSSION

Many attempts have been made at enhancing wheat transformation using immature embryos. However, quite often, those effects resulted in unintended complexity for conducting transformation. For example, some research has suggested pre-culturing the embryos to increase transformation efficiency (Kumar et al., 2019). Others have used surfactants, such as Silwet L-77, to increase T-DNA delivery (Hayta et al., 2019). A recently reported protocol requires shaking the embryos during infection (Raman et al., 2022). Refrigeration while centrifuging is also very common (Ishida et al., 2015; Hayta et al., 2019). Finally, most methods require the excision of the embryo axis following co-cultivation (Ishida et al., 2015; Hayta et al., 2019; Wang et al., 2022). The QuickWheat transformation system described herein does not require such complexity as there is no pre-culturing of embryos, no surfactants, shaking, or embryo axis excision, and centrifugation does not require refrigeration.

### Non-Excision Wheat Transformation Method

The *Agrobacterium*-mediated QuickWheat transformation system uses a thymidine auxotrophic *Agrobacterium* strain (LBA4404 Thy-) harboring the ternary vector system with accessory plasmid pPHP71539 (pVir) for T-DNA delivery as described previously for maize (Anand et al., 2018) and sorghum (Che et al., 2018; Che et al., 2022) transformation. This T-DNA delivery system was used for developing both non-excision and excision QuickWheat transformation methods as described in the Methods. The non-excision binary T-DNA vector carried *ZmWus2* and *ZmBbm* (*Zm-Axig1_pro_* or *Zm-PTLP_pro_:ZmWus2* and *Zm-PLTP_pro_:ZmBbm*) as morphogenic genes, the gene of interest (GOI) as trait genes (e.g., genome editing components including Cas9 and gRNAs) and *NPTII* (*Ubi_pro_:NPTII*) or *moPAT* (*Ubi_pro_:moPAT*) as a selectable marker (**Figure 2A**, **B** and **C**). In one of the experiments, as shown in **Table 3**, 162 spring wheat SBC0456D immature embryos isolated from triplicated biological replications were infected with *Agrobacterium* carrying the construct as illustrated in **Figure 2A**. The isolated immature embryos were pretreated by centrifugation at room temperature (refrigerated centrifugation is not required) to increase *Agrobacterium*-mediated T-DNA gene delivery. After *Agrobacterium* infection and co-cultivation, the immature embryos were then sub-cultured on culture medium #300 (**Table 2**) to induce callus and rapid somatic embryo formation (**Figure 1F**, and **G**). Shoot regeneration and rooting was induced in the somatic embryos on culture medium #400 (**Table 2**) with selection (150 mg/L G418) to generate transgenic plantlets before transfer to soil. Due to the rapid somatic embryogenesis induced by ZmWus2 and ZmBbm, embryonic axis excision and the prolonged dual-selection callus formation, steps which are reported to be essential for conventional transformation methods (Ishida et al., 2015; Hayta et al., 2019) (**Figure 3A**), were unnecessary for this QuickWheat transformation protocol. Furthermore, the prerequisite of cytokinin on shoot regeneration medium was no longer necessary for shoot induction and elongation. The total time from inoculation of immature embryos to transplantation of a fully developed transgenic plants to the greenhouse was reduced to 53 days (**Figure 3B**) compared to around 80 days using the conventional method (**Figure 3A**). As shown in **Table 3**, out of 162 infected embryos, 122 had at least one regenerated shoot develop. Following shoot and root regeneration and elongation, a total of 245 plantlets were isolated from 122 embryos (Multiple shoots can be developed and isolated from individual embryos, from which less than 20% of the events were dependent events as demonstrated below). Therefore, the 75% ‘independent-transformation efficiency’ was calculated as the total number of embryos with at least one regenerated shoot over the total number of inoculated embryos and the 151% ‘overall-transformation efficiency’ was calculated as the total number isolated plantlets over the total number of inoculated embryos. For the 245 plants analyzed, 25% of the events were single-copy quality event (QE) (single copy and backbone-free) and the rest of 75% were non-QE events in which 5% were escapes (**Table 3**). To test the flexibility of the non-excision QuickWheat transformation method established, we further conducted transformation in different genotypic backgrounds using different selectable markers. As shown in **Table 3**, an average of 57% independent-transformation efficiencies was achieved in Fielder using *NPTII* as selectable marker (**Figure 2B**) based on seven biological replications. The genotype Chinese Spring reached 94% overall-transformation efficiency using *moPAT* as selectable marker (**Figure 2C**) based on five biological replications.

**TABLE 3.**
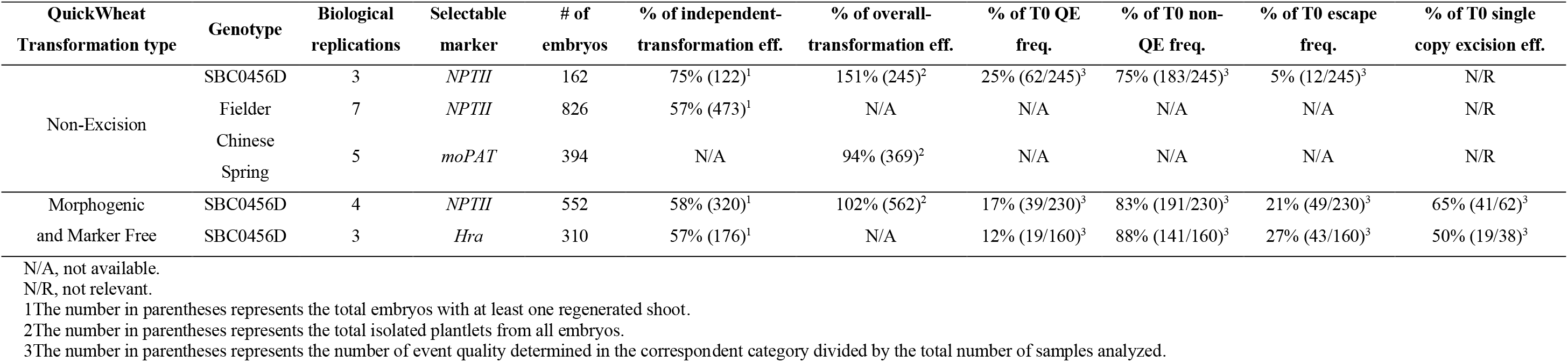
Transformation efficiency and event quality frequency of QuickWheat transformation system

Pleiotropic effects have been reported in transgenic maize and sorghum plants carrying *ZmBbm* and *ZmWus2* genes (Lowe et al., 2016; Che et al., 2022). To further evaluate the impact of morphogenic gene expression on plant growth and fertility in wheat, transgenic T0 plants transformed with a non-excision construct (**Figure 2A**) were sent to the greenhouse and grown to fertility. As shown in **Figure 1N**, no obvious pleiotropic effects were observed for any transgenic SBC0456D plants (**Figure 1N**) compared to the wild-type (**Figure 1M**) and all the transgenic events were fertile regardless of the copy number of the transgene.

To determine whether multiple plants from one embryo were independent or dependent (clonal), a separate transformation experiment was initiated using the same non-excision construct in **Figure 2A**. We identified 35 embryos that generated multiple transgenic shoots. A total of 100 individual plantlets were isolated from those 35 embryos and leaf samples were collected individually to determine the dependency of the events isolated from the same embryos. The independency or dependency of the events was determined based on 13 assays as described in the Methods and as shown in **Supplementary Table 1** and **Supplementary Figure 2**. Nine assays were designed to determine T-DNA integration. Of these nine assays, three revealed copy number and six detected presence/absence of the T-DNA elements. Additionally, four assays were used for testing presence/absence of T-DNA backbone integrations. A call of “independent” or “presumably dependent” was made for each individual plantlet depending on the results from other plants regenerated from the same embryo. A “presumably dependent” call was made if multiple events isolated from the same embryo had all 13 identical assays. If at least one assay out of the 13 assays differed, they were defined as “independent” events. As shown in **Supplementary Table 1**, 84 plantlets isolated from 28 embryos showed at least one different assay when compared to other plants from the same embryo, suggesting that these plantlets were independent transformation events. The remaining 16 plantlets isolated from seven embryos had 13 identical assay results. Therefore, the transformation dependency of those 16 events were uncertain (presumably dependent) pending determination by more specific analyses, such as Southern-by-Sequencing analysis. As such, the current data suggests that there is no more than 16% chance (16 plants out of 100 sampled) that multiple plantlets isolated from a single embryo were dependent events. Furthermore, based on the 100 plantlets analyzed from the 35 embryos, there were 15 single-copy QEs, of which only two were from the same embryo. This QE pair was presumably dependent due to identical assay results (**Supplementary Table 1**), suggesting that it is even rarer, 7% (1 out of 15 QEs), chance of isolating dependent QEs from the same embryo. Therefore, isolating multiple events from one embryo for QE identification is a feasible approach to increase the frequency of QEs and reduce workload for conducting transformation. This approach should be even more attractive for generating genome editing events because transgenes can be segregated out from the edited allele(s) in the subsequent generations, and the event quality and dependency are, therefore, no longer a concern.

### Excision Wheat Transformation Method

Selectable marker genes, such as antibiotic or herbicide resistance genes, are often used for plant transformation to select transgenic events (Breyer et al., 2014). The presence of marker genes in transgenic plants, however, often provides no advantage after transformation and may raise biosafety concerns for commercial release (Breyer et al., 2014). In addition, morphogenic genes (*ZmWus2* and *ZmBbm*) used herein for achieving highly efficient transformation are transcription factors that affect many aspects of plant development, such as cell division, differentiation, proliferation and reproduction, etc. (Gordon-Kamm et al., 2019). Although no pleiotropic effects were observed in the transgenic wheat plants using the non-excision transformation method as described above, continued expression of morphogenic genes could interfere with downstream trait gene functional analysis and characterization. Therefore, we chose to develop a more advanced excision system to enable morphogenic- and marker-gene-free transformation. As illustrated in **Figure 2D** and **E**, the morphogenic- and marker-gene-free binary vector contains *ZmWus2* (*Zm-PLTP_pro_:ZmWus2*), a heat shock inducible *moCRE* gene (*Zm-Hsp17.7_pro_:moCRE*) and a selectable marker (*UBI_pro_:NPTII*) or (*Sb-Als_pro_:ZmHra*) flanked by repeated *loxP* sites. The GOI was inserted outside of *loxP* sites.

Following *Agrobacterium* infection using the strain carrying the construct as illustrated in **Figure 2D** and co-cultivation as described in the Methods, SBC0456D immature embryos were then sub-cultured on culture medium #300 (**Table 2**) with selection (100 mg/L G418) for three weeks to induce somatic embryo formation (**Figure 3B**). The intent of this step was to maximize transgenic somatic embryo formation, but at the same time minimize the proliferation of non-transgenic tissue to enrich for transgenic somatic embryos survival and germination during regeneration and elongation without selection. After three-week long stringent selection, the excision of the *ZmWus2/moCRE/NPTII* cassette (**Figure 2D**) was accomplished by inducing *moCRE* expression upon heat treatment at 45°C and 70% humidity for two hours (**Figure 3B**). After heat-shock treatment, embryos were transferred to the regeneration culture medium #400 (**Table 2**) without selection for 4-5 weeks to induce shoot regeneration, elongation, and rooting. The total time from inoculation of immature embryos to transplantation of fully developed transgenic plants to the greenhouse took 57 to 64 days. As shown in **Table 3**, the three-week long selection pressure during somatic embryo formation was stringent and produced no more than 21% escapes based on four biological replications. The heat-shock-induced *moCRE/LoxP*-mediated excision system was highly efficient as well, with more than 65% of the single-copy events showing complete *ZmWus2/moCRE/NPTII* cassette excision (**Table 3**). Although both the independent-transformation efficiency (58%) and overall-transformation efficiency (102%) were lower compared to the non-excision transformation method, 75% and 151% respectively, the overall QE frequency (single copy, backbone-free and *ZmWus2/moCRE/NPTII* cassette-free) was as high as 17% (**Table 3**), just slightly lower than 25% QE frequency (single copy and backbone-free only) for the non-excision transformation method. The high QE frequency of this morphogenic- and marker-gene-free method most likely resulted from of highly efficient excision efficiency in single-copy events (65%) and moderate escape frequency (21%).

To broaden the application of this method, we further tested the morphogenic- and marker-gene-free method using *Hra* as the selectable marker based on three biological replications (**Figure 2E**) and achieved similar independent-transformation efficiency (57%), QE frequency (12%) and excision efficiency (50%) compared to *NPTII* (**Figure 2D and Table 3**).

In summary, we have developed the rapid, robust, and highly efficient QuickWheat transformation system for broad applications to make the transformation process less labor-intensive and more cost-effective. In addition, the QuickWheat methods developed are flexible for multiple selectable marker systems, highly reproducible, high-throughput and have been routinely used in Corteva transformation production setting in three wheat genotypes for the past few years to generate thousands of transgenic events, including CRISPR/Cas-mediated genome editing events (editing data was not presented in this manuscript and will be published separately). We believe that QuickWheat transformation can easily be implemented by most laboratories and will have an immediate and far-reaching impact on wheat research.

### Materials Availability

Novel biological materials described in this publication may be available to the academic community and other not-for-profit institutions solely for non-commercial research purposes upon acceptance and signing of a material transfer agreement between the author’s institution and the requestor. In some cases, such materials may contain genetic elements described in the manuscript that were obtained from a third party(s) (e.g., *Cas9*) and the authors may not be able to provide materials including third party genetic elements to the requestor because of certain third-party contractual restrictions placed on the author’s institution. In such cases, the requester will be required to obtain such materials directly from the third party. The author’s and authors’ institution do not make any express or implied permission(s) to the requester to make, use, sell, offer for sale, or import third party proprietary materials. Obtaining any such permission(s) will be the sole responsibility of the requestor. Corteva Agriscience™ proprietary plant germplasm and any transgenic material will not be made available except at the discretion of the owner and then only in accordance with all applicable governmental regulations.

## Acknowledgements

We thank the super binary vector construction team from Corteva Agriscience for their support with vector construction, and PCR analysis characterization group from Corteva Agriscience for event quality analysis and environmental control group from Corteva Agriscience for wheat planting in the greenhouse.

## Author Contributions

All authors are involved in designing research, analyzing the data, and writing the paper. K.J. conducted wheat transformation.

## SUPPLEMENTARY MATERIAL

**SUPPLEMENTARY FIGURE 1** Tools used for wheat transformation. (**A**) Tabletop centrifuge. (**B**) Embryo isolation tool.

**SUPPLEMENTARY FIGURE 2** Schematic representation of the qPCR assays for event quality and dependency analysis. Event quality and dependency were analyzed by 13 assays (in green).

**SUPPLEMENTARY TABLE 1** Event dependency determination

