## Supplementary material for "Rapid and Highly Efficient Morphogenic Gene-Mediated Hexaploid Wheat Transformation"

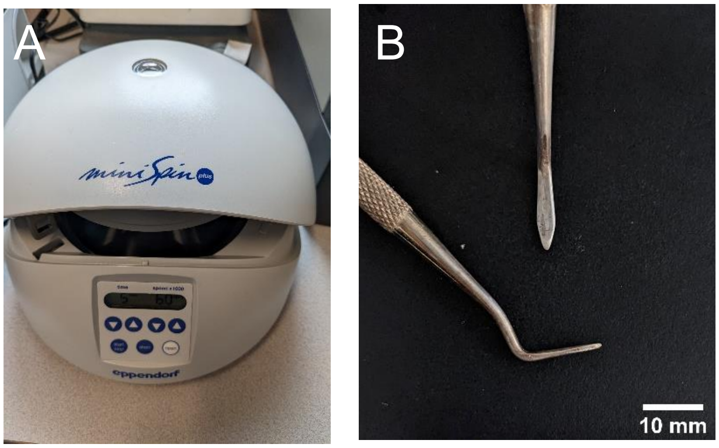


**SUPPLEMENTARY FIGURE 1** Tools used for wheat transformation. (**A**) Tabletop centrifuge. (**B**) Embryo isolation tool.

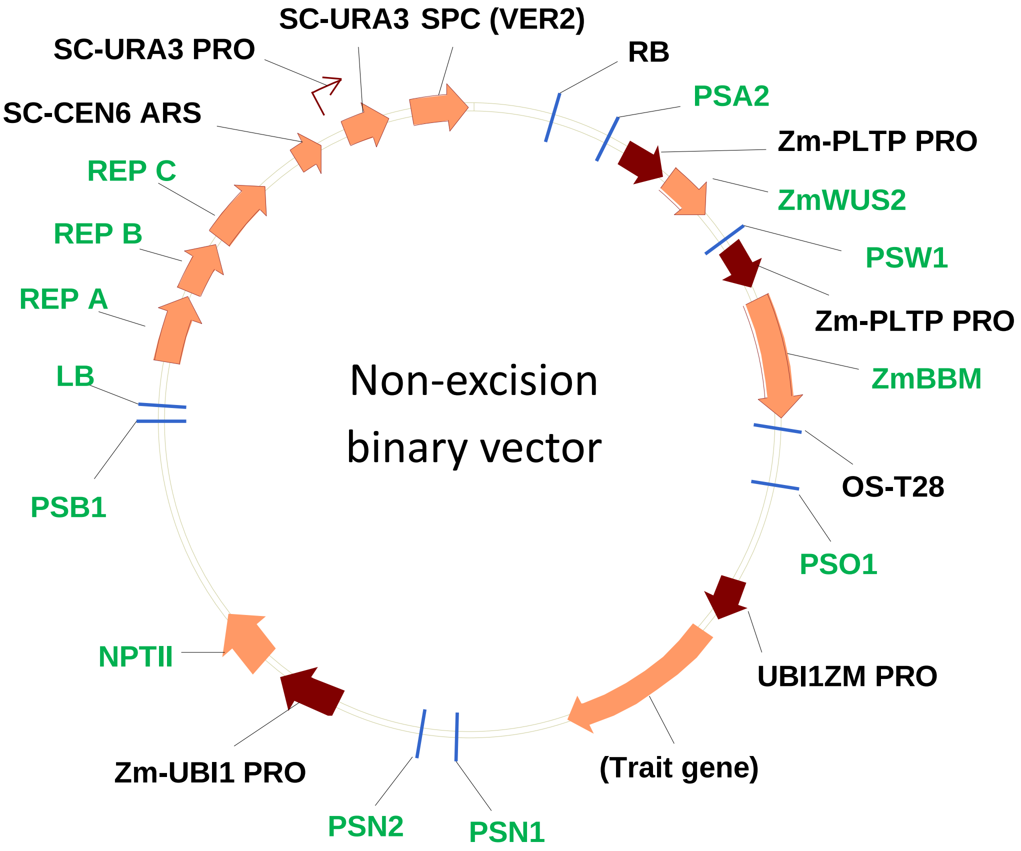


**SUPPLEMENTARY FIGURE 2** Schematic representation of the qPCR assays for event quality and dependency analysis. Event quality and dependency were analyzed by 13 assays (in green).

**SUPPLEMENTARY TABLE 1** Event dependency determination

| Embryo | Individual Plantlet | QE Call | Event Dependency Call | PSA2 | NPTII | PSB1 | ZmWUS2 | PSW1 | ZmBBM | PSO1 | PSN1 | PSN2 | LB | REP A | REP B | REP C |
|---|---|---|---|---|---|---|---|---|---|---|---|---|---|---|---|---|
| 1 | 1a | Non-QE | Independent | NULL | 1 | 1 | NEGATIVE | NEGATIVE | NEGATIVE | NEGATIVE | NEGATIVE | NEGATIVE | NEGATIVE | NEGATIVE | NEGATIVE | NEGATIVE |
| | 1b | QE | Independent | 1 | 1 | 1 | POSITIVE | POSITIVE | POSITIVE | POSITIVE | POSITIVE | POSITIVE | NEGATIVE | NEGATIVE | NEGATIVE | NEGATIVE |
| | 1c | Non-QE | Independent | 2 | 2 | 3 | POSITIVE | POSITIVE | POSITIVE | POSITIVE | POSITIVE | POSITIVE | NEGATIVE | NEGATIVE | NEGATIVE | NEGATIVE |
| 2 | 2a | Non-QE | Independent | 1 | 2 | 3 | POSITIVE | POSITIVE | POSITIVE | POSITIVE | POSITIVE | POSITIVE | NEGATIVE | NEGATIVE | NEGATIVE | NEGATIVE |
| | 2b | Non-QE | Independent | 3 | 1 | 1 | POSITIVE | POSITIVE | POSITIVE | POSITIVE | POSITIVE | POSITIVE | NEGATIVE | NEGATIVE | NEGATIVE | NEGATIVE |
| | 2c | QE | Independent | 1 | 1 | 1 | POSITIVE | POSITIVE | POSITIVE | POSITIVE | POSITIVE | POSITIVE | NEGATIVE | NEGATIVE | NEGATIVE | NEGATIVE |
| | 2d | Non-QE | Independent | NULL | 1 | 1 | NEGATIVE | NEGATIVE | NEGATIVE | NEGATIVE | POSITIVE | POSITIVE | NEGATIVE | NEGATIVE | NEGATIVE | NEGATIVE |
| 3 | 3a | Non-QE | Independent | 4 | 3 | 1 | POSITIVE | POSITIVE | POSITIVE | POSITIVE | POSITIVE | POSITIVE | NEGATIVE | NEGATIVE | NEGATIVE | NEGATIVE |
| | 3b | Non-QE | Independent | 2 | 2 | 1 | POSITIVE | POSITIVE | POSITIVE | POSITIVE | POSITIVE | POSITIVE | NEGATIVE | NEGATIVE | NEGATIVE | NEGATIVE |
| | 3c | Non-QE | Independent | 2 | 1 | 1 | POSITIVE | POSITIVE | POSITIVE | POSITIVE | POSITIVE | POSITIVE | NEGATIVE | NEGATIVE | NEGATIVE | NEGATIVE |
| 4 | 4a | Non-QE | Independent | 1 | 1 | 1 | POSITIVE | POSITIVE | POSITIVE | POSITIVE | POSITIVE | POSITIVE | POSITIVE | POSITIVE | POSITIVE | POSITIVE |
| | 4b | Non-QE | Independent | 4 | 3 | 4 | POSITIVE | POSITIVE | POSITIVE | POSITIVE | POSITIVE | POSITIVE | NEGATIVE | NEGATIVE | NEGATIVE | NEGATIVE |
| | 4c | Non-QE | Independent | 1 | 1 | 2 | POSITIVE | POSITIVE | POSITIVE | POSITIVE | POSITIVE | POSITIVE | NEGATIVE | NEGATIVE | NEGATIVE | NULL |
| | 4d | Non-QE | Independent | 2 | 1 | 1 | POSITIVE | POSITIVE | POSITIVE | POSITIVE | POSITIVE | POSITIVE | NEGATIVE | NEGATIVE | NEGATIVE | NEGATIVE |
| | 4e | Non-QE | Independent | 4 | 2 | 2 | POSITIVE | POSITIVE | POSITIVE | POSITIVE | POSITIVE | POSITIVE | NEGATIVE | NEGATIVE | NEGATIVE | NEGATIVE |
| | 4f | QE | Independent | 1 | 1 | 1 | POSITIVE | POSITIVE | POSITIVE | POSITIVE | POSITIVE | POSITIVE | NEGATIVE | NEGATIVE | NEGATIVE | NEGATIVE |
| 5 | 5a | Non-QE | Independent | 2 | 1 | 1 | POSITIVE | POSITIVE | POSITIVE | POSITIVE | POSITIVE | POSITIVE | NEGATIVE | NEGATIVE | NEGATIVE | NEGATIVE |
| | 5b | QE | Independent | 1 | 1 | 1 | POSITIVE | POSITIVE | POSITIVE | POSITIVE | POSITIVE | POSITIVE | NEGATIVE | NEGATIVE | NEGATIVE | NEGATIVE |
| 6 | 6a | Non-QE | Independent | 2 | 1 | 1 | POSITIVE | POSITIVE | POSITIVE | POSITIVE | POSITIVE | POSITIVE | NEGATIVE | NEGATIVE | NEGATIVE | NEGATIVE |
| | 6b | QE | Independent | 1 | 1 | 1 | POSITIVE | POSITIVE | POSITIVE | POSITIVE | POSITIVE | POSITIVE | NEGATIVE | NEGATIVE | NEGATIVE | NEGATIVE |
| 7 | 7a | Non-QE | Independent | 2 | 2 | 3 | POSITIVE | POSITIVE | POSITIVE | POSITIVE | POSITIVE | POSITIVE | NEGATIVE | NEGATIVE | NEGATIVE | NEGATIVE |
| | 7b | Non-QE | Independent | 1 | 1 | 3 | POSITIVE | POSITIVE | POSITIVE | POSITIVE | POSITIVE | POSITIVE | NEGATIVE | NEGATIVE | NEGATIVE | NEGATIVE |
| 8 | 8a | Non-QE | Independent | 1 | 2 | 2 | POSITIVE | POSITIVE | POSITIVE | POSITIVE | POSITIVE | POSITIVE | NEGATIVE | NEGATIVE | NEGATIVE | NEGATIVE |
| | 8b | Non-QE | Independent | 3 | 2 | 2 | POSITIVE | POSITIVE | POSITIVE | POSITIVE | POSITIVE | POSITIVE | NEGATIVE | NEGATIVE | NEGATIVE | NEGATIVE |
| 9 | 9a | Non-QE | Independent | 2 | 2 | 2 | POSITIVE | POSITIVE | POSITIVE | POSITIVE | POSITIVE | POSITIVE | NEGATIVE | NEGATIVE | NEGATIVE | NEGATIVE |
| | 9b | QE | Independent | 1 | 1 | 1 | POSITIVE | POSITIVE | POSITIVE | POSITIVE | POSITIVE | POSITIVE | NEGATIVE | NEGATIVE | NEGATIVE | NEGATIVE |
| 10 | 10a | QE | Independent | 1 | 1 | 1 | POSITIVE | POSITIVE | POSITIVE | POSITIVE | POSITIVE | POSITIVE | NEGATIVE | NEGATIVE | NEGATIVE | NEGATIVE |
| | 10b | Non-QE | Independent | NULL | 1 | 1 | POSITIVE | POSITIVE | POSITIVE | POSITIVE | POSITIVE | POSITIVE | NEGATIVE | NEGATIVE | NEGATIVE | NEGATIVE |
| 11 | 11a | Non-QE | Independent | 1 | 1 | 1 | NEGATIVE | NEGATIVE | NEGATIVE | NEGATIVE | POSITIVE | POSITIVE | NEGATIVE | NEGATIVE | NEGATIVE | NEGATIVE |
| | 11b | Non-QE | Independent | 4 | 2 | 2 | POSITIVE | POSITIVE | POSITIVE | POSITIVE | POSITIVE | POSITIVE | NEGATIVE | NEGATIVE | NEGATIVE | NEGATIVE |
| 12 | 12a | Non-QE | Independent | 3 | 2 | 3 | POSITIVE | POSITIVE | POSITIVE | POSITIVE | POSITIVE | POSITIVE | NEGATIVE | NEGATIVE | NEGATIVE | NEGATIVE |
| | 12b | Non-QE | Independent | 2 | 1 | 2 | POSITIVE | POSITIVE | POSITIVE | POSITIVE | POSITIVE | POSITIVE | NEGATIVE | NEGATIVE | NEGATIVE | NEGATIVE |
| 13 | 13a | Non-QE | Independent | 2 | 3 | 3 | POSITIVE | POSITIVE | POSITIVE | POSITIVE | POSITIVE | POSITIVE | NEGATIVE | NEGATIVE | NEGATIVE | NEGATIVE |
| | 13b | Non-QE | Independent | 3 | 3 | 2 | POSITIVE | POSITIVE | POSITIVE | POSITIVE | POSITIVE | POSITIVE | NEGATIVE | NEGATIVE | NEGATIVE | NEGATIVE |
| | 13c | Non-QE | Independent | 4 | 2 | 2 | POSITIVE | POSITIVE | POSITIVE | POSITIVE | POSITIVE | POSITIVE | POSITIVE | POSITIVE | POSITIVE | POSITIVE |
| 14 | 14a | Non-QE | Independent | 1 | 2 | 2 | POSITIVE | POSITIVE | POSITIVE | POSITIVE | POSITIVE | POSITIVE | POSITIVE | POSITIVE | POSITIVE | POSITIVE |
| | 14b | QE | Independent | 1 | 1 | 1 | POSITIVE | POSITIVE | POSITIVE | POSITIVE | POSITIVE | POSITIVE | NEGATIVE | NEGATIVE | NEGATIVE | NEGATIVE |
| 15 | 15a | Non-QE | Independent | 3 | 3 | 3 | POSITIVE | POSITIVE | POSITIVE | POSITIVE | POSITIVE | POSITIVE | NEGATIVE | NEGATIVE | NEGATIVE | NEGATIVE |
| | 15b | Non-QE | Independent | 3 | 2 | 1 | POSITIVE | POSITIVE | POSITIVE | POSITIVE | POSITIVE | POSITIVE | NEGATIVE | NEGATIVE | NEGATIVE | NEGATIVE |
| 16 | 16a | Non-QE | Independent | 3 | 1 | 1 | POSITIVE | POSITIVE | POSITIVE | POSITIVE | POSITIVE | POSITIVE | NEGATIVE | NEGATIVE | NEGATIVE | NEGATIVE |
| | 16b | QE | Independent | 1 | 1 | 1 | POSITIVE | POSITIVE | POSITIVE | POSITIVE | POSITIVE | POSITIVE | NEGATIVE | NEGATIVE | NEGATIVE | NEGATIVE |
| | 16c | Non-QE | Independent | 2 | 1 | 2 | POSITIVE | POSITIVE | POSITIVE | POSITIVE | POSITIVE | POSITIVE | NEGATIVE | NEGATIVE | NEGATIVE | NEGATIVE |
| 17 | 17a | Non-QE | Independent | 1 | 1 | 2 | POSITIVE | POSITIVE | POSITIVE | POSITIVE | POSITIVE | POSITIVE | NEGATIVE | NEGATIVE | NEGATIVE | NEGATIVE |
| | 17b | Non-QE | Independent | 1 | 1 | NULL | POSITIVE | POSITIVE | POSITIVE | POSITIVE | POSITIVE | POSITIVE | NEGATIVE | NEGATIVE | NEGATIVE | NEGATIVE |
| 18 | 18a | Non-QE | Independent | 2 | 1 | 1 | POSITIVE | POSITIVE | POSITIVE | POSITIVE | POSITIVE | POSITIVE | NEGATIVE | NEGATIVE | NEGATIVE | NEGATIVE |
| | 18b | QE | Independent | 1 | 1 | 1 | POSITIVE | POSITIVE | POSITIVE | POSITIVE | POSITIVE | POSITIVE | NEGATIVE | NEGATIVE | NEGATIVE | NEGATIVE |
| 19 | 19a | Non-QE | Independent | 4 | 2 | 2 | POSITIVE | POSITIVE | POSITIVE | POSITIVE | POSITIVE | POSITIVE | NEGATIVE | NEGATIVE | NEGATIVE | NEGATIVE |
| | 19b | Non-QE | Independent | 2 | 1 | 1 | POSITIVE | POSITIVE | POSITIVE | POSITIVE | POSITIVE | POSITIVE | NEGATIVE | NEGATIVE | NEGATIVE | NEGATIVE |
| | 19c | QE | Independent | 1 | 1 | 1 | POSITIVE | POSITIVE | POSITIVE | POSITIVE | POSITIVE | POSITIVE | NEGATIVE | NEGATIVE | NEGATIVE | NEGATIVE |
| | 19d | Non-QE | Independent | 4 | 3 | 4 | POSITIVE | POSITIVE | POSITIVE | POSITIVE | POSITIVE | POSITIVE | NEGATIVE | NEGATIVE | NEGATIVE | NEGATIVE |
| 20 | 20a | Non-QE | Independent | 2 | 1 | 1 | POSITIVE | POSITIVE | POSITIVE | POSITIVE | POSITIVE | POSITIVE | NEGATIVE | NEGATIVE | NEGATIVE | NEGATIVE |
| | 20b | Non-QE | Independent | 4 | 1 | 1 | POSITIVE | POSITIVE | POSITIVE | POSITIVE | POSITIVE | POSITIVE | NEGATIVE | NEGATIVE | NEGATIVE | NEGATIVE |
| 21 | 21a | Non-QE | Independent | 1 | 1 | 2 | POSITIVE | POSITIVE | POSITIVE | POSITIVE | POSITIVE | POSITIVE | NEGATIVE | NEGATIVE | NEGATIVE | NEGATIVE |
| | 21b | Non-QE | Independent | >4COPY | 4 | 4 | POSITIVE | POSITIVE | POSITIVE | POSITIVE | POSITIVE | POSITIVE | POSITIVE | POSITIVE | POSITIVE | POSITIVE |
| 22 | 22a | Non-QE | Independent | 4 | 3 | >4COPY | POSITIVE | POSITIVE | POSITIVE | POSITIVE | POSITIVE | POSITIVE | NEGATIVE | NEGATIVE | NEGATIVE | NEGATIVE |
| | 22b | Non-QE | Independent | 1 | 1 | 2 | POSITIVE | POSITIVE | POSITIVE | POSITIVE | POSITIVE | POSITIVE | NEGATIVE | NEGATIVE | NEGATIVE | NEGATIVE |
| | 22c | Non-QE | Independent | 4 | 2 | 2 | POSITIVE | POSITIVE | POSITIVE | POSITIVE | POSITIVE | POSITIVE | NEGATIVE | NEGATIVE | NEGATIVE | NEGATIVE |
| 23 | 23a | Non-QE | Independent | 2 | 2 | 2 | POSITIVE | POSITIVE | POSITIVE | POSITIVE | POSITIVE | POSITIVE | NEGATIVE | NEGATIVE | NEGATIVE | NEGATIVE |
| | 23b | QE | Independent | 1 | 1 | 1 | POSITIVE | POSITIVE | POSITIVE | POSITIVE | POSITIVE | POSITIVE | NEGATIVE | NEGATIVE | NEGATIVE | NEGATIVE |
| 24 | 24a | QE | Independent | 1 | 1 | 1 | POSITIVE | POSITIVE | POSITIVE | POSITIVE | POSITIVE | POSITIVE | NEGATIVE | NEGATIVE | NEGATIVE | NEGATIVE |
| | 24b | Non-QE | Independent | 2 | 1 | 3 | POSITIVE | POSITIVE | POSITIVE | POSITIVE | POSITIVE | POSITIVE | NEGATIVE | NEGATIVE | NEGATIVE | NEGATIVE |
| | 24c | Non-QE | Independent | 4 | 2 | 2 | POSITIVE | POSITIVE | POSITIVE | POSITIVE | POSITIVE | POSITIVE | NEGATIVE | NEGATIVE | NEGATIVE | NEGATIVE |
| 25 | 25a | Non-QE | Independent | 2 | 1 | 1 | POSITIVE | POSITIVE | POSITIVE | POSITIVE | POSITIVE | POSITIVE | NEGATIVE | NEGATIVE | NEGATIVE | NEGATIVE |
| | 25b | Non-QE | Independent | 4 | 2 | 1 | POSITIVE | POSITIVE | POSITIVE | POSITIVE | POSITIVE | POSITIVE | POSITIVE | NEGATIVE | NEGATIVE | NEGATIVE |
| | 25c | Non-QE | Independent | 1 | 2 | 3 | POSITIVE | POSITIVE | POSITIVE | POSITIVE | POSITIVE | POSITIVE | NEGATIVE | NEGATIVE | NEGATIVE | NEGATIVE |
| 26 | 26a | Non-QE | Independent | 4 | 2 | 1 | POSITIVE | POSITIVE | POSITIVE | POSITIVE | POSITIVE | POSITIVE | NEGATIVE | POSITIVE | NEGATIVE | NEGATIVE |
| | 26b | Non-QE | Independent | 2 | 1 | NULL | POSITIVE | POSITIVE | POSITIVE | POSITIVE | POSITIVE | POSITIVE | NEGATIVE | NEGATIVE | NEGATIVE | NEGATIVE |
| 27 | 27a | Non-QE | Independent | 1 | 1 | NULL | POSITIVE | POSITIVE | POSITIVE | POSITIVE | POSITIVE | POSITIVE | NEGATIVE | NEGATIVE | NEGATIVE | NEGATIVE |
| | 27b | Non-QE | Independent | 2 | 1 | 1 | POSITIVE | POSITIVE | POSITIVE | POSITIVE | POSITIVE | POSITIVE | POSITIVE | POSITIVE | POSITIVE | POSITIVE |
| 28 | 28a | Non-QE | Independent | 2 | 1 | NULL | POSITIVE | POSITIVE | POSITIVE | POSITIVE | POSITIVE | POSITIVE | NEGATIVE | NEGATIVE | NEGATIVE | NEGATIVE |
| | 28b | Non-QE | Independent | 2 | 1 | 1 | POSITIVE | POSITIVE | POSITIVE | POSITIVE | POSITIVE | POSITIVE | NEGATIVE | NEGATIVE | NEGATIVE | NEGATIVE |
| | 28c | Non-QE | Independent | 3 | 3 | 2 | POSITIVE | POSITIVE | POSITIVE | POSITIVE | POSITIVE | POSITIVE | NEGATIVE | NEGATIVE | NEGATIVE | NEGATIVE |
| | 28d | Non-QE | Independent | 1 | 1 | NULL | POSITIVE | POSITIVE | POSITIVE | POSITIVE | POSITIVE | POSITIVE | NEGATIVE | NEGATIVE | NEGATIVE | NEGATIVE |
| 29 | 29a | Non-QE | Presumably Dependent | 4 | 4 | 4 | POSITIVE | POSITIVE | POSITIVE | POSITIVE | POSITIVE | POSITIVE | NEGATIVE | NEGATIVE | NEGATIVE | NEGATIVE |
| | 29b | Non-QE | | 4 | 4 | 4 | POSITIVE | POSITIVE | POSITIVE | POSITIVE | POSITIVE | POSITIVE | NEGATIVE | NEGATIVE | NEGATIVE | NEGATIVE |
| | 29c | Non-QE | Independent | 1 | 1 | 2 | POSITIVE | POSITIVE | POSITIVE | POSITIVE | POSITIVE | POSITIVE | NEGATIVE | NEGATIVE | NEGATIVE | NEGATIVE |
| 30 | 30a | Non-QE | Presumably Dependent | 3 | 2 | 2 | POSITIVE | POSITIVE | POSITIVE | POSITIVE | POSITIVE | POSITIVE | NEGATIVE | NEGATIVE | NEGATIVE | NEGATIVE |
| | 30b | Non-QE | | 3 | 2 | 2 | POSITIVE | POSITIVE | POSITIVE | POSITIVE | POSITIVE | POSITIVE | NEGATIVE | NEGATIVE | NEGATIVE | NEGATIVE |
| | 30c | QE | Presumably Dependent | 1 | 1 | 1 | POSITIVE | POSITIVE | POSITIVE | POSITIVE | POSITIVE | POSITIVE | NEGATIVE | NEGATIVE | NEGATIVE | NEGATIVE |
| | 30d | QE | | 1 | 1 | 1 | POSITIVE | POSITIVE | POSITIVE | POSITIVE | POSITIVE | POSITIVE | NEGATIVE | NEGATIVE | NEGATIVE | NEGATIVE |
| | 30e | Non-QE | Independent | 3 | 2 | 1 | POSITIVE | POSITIVE | POSITIVE | POSITIVE | POSITIVE | POSITIVE | NEGATIVE | NEGATIVE | NEGATIVE | NEGATIVE |
| | 30f | Non-QE | Independent | 2 | 1 | 1 | POSITIVE | POSITIVE | POSITIVE | POSITIVE | POSITIVE | POSITIVE | NEGATIVE | NEGATIVE | NEGATIVE | NEGATIVE |
| | 30g | Non-QE | Independent | 3 | 1 | 1 | POSITIVE | POSITIVE | POSITIVE | POSITIVE | POSITIVE | POSITIVE | NEGATIVE | NEGATIVE | NEGATIVE | NEGATIVE |
| 31 | 31a | Non-QE | Presumably Dependent | 2 | 1 | 1 | POSITIVE | POSITIVE | POSITIVE | POSITIVE | POSITIVE | POSITIVE | NEGATIVE | NEGATIVE | NEGATIVE | NEGATIVE |
| | 31b | Non-QE | | 2 | 1 | 1 | POSITIVE | POSITIVE | POSITIVE | POSITIVE | POSITIVE | POSITIVE | NEGATIVE | NEGATIVE | NEGATIVE | NEGATIVE |
| | 31c | Non-QE | Independent | 2 | 1 | 2 | POSITIVE | POSITIVE | POSITIVE | POSITIVE | POSITIVE | POSITIVE | NEGATIVE | NEGATIVE | NEGATIVE | NEGATIVE |
| | 31d | Non-QE | Independent | 1 | 1 | WEAK POSITIVE | POSITIVE | POSITIVE | POSITIVE | POSITIVE | POSITIVE | POSITIVE | NEGATIVE | NEGATIVE | NEGATIVE | NEGATIVE |
| | 31e | Non-QE | Independent | >4COPY | >4COPY | 3 | POSITIVE | POSITIVE | POSITIVE | POSITIVE | POSITIVE | POSITIVE | POSITIVE | POSITIVE | POSITIVE | POSITIVE |
| 32 | 32a | Non-QE | Presumably Dependent | 1 | 1 | 2 | POSITIVE | POSITIVE | POSITIVE | POSITIVE | POSITIVE | POSITIVE | NEGATIVE | NEGATIVE | NEGATIVE | NEGATIVE |
| | 32b | Non-QE | | 1 | 1 | 2 | POSITIVE | POSITIVE | POSITIVE | POSITIVE | POSITIVE | POSITIVE | NEGATIVE | NEGATIVE | NEGATIVE | NEGATIVE |
| | 32c | QE | Independent | 1 | 1 | 1 | POSITIVE | POSITIVE | POSITIVE | POSITIVE | POSITIVE | POSITIVE | NEGATIVE | NEGATIVE | NEGATIVE | NEGATIVE |
| | 32d | Non-QE | Independent | 1 | 1 | 3 | POSITIVE | POSITIVE | NEGATIVE | NEGATIVE | POSITIVE | POSITIVE | NEGATIVE | NEGATIVE | NEGATIVE | NEGATIVE |
| 33 | 33a | Non-QE | Presumably Dependent | 2 | 1 | 1 | POSITIVE | POSITIVE | POSITIVE | POSITIVE | POSITIVE | POSITIVE | NEGATIVE | NEGATIVE | NEGATIVE | NEGATIVE |
| | 33b | Non-QE | | 2 | 1 | 1 | POSITIVE | POSITIVE | POSITIVE | POSITIVE | POSITIVE | POSITIVE | NEGATIVE | NEGATIVE | NEGATIVE | NEGATIVE |
| 34 | 34a | Non-QE | Presumably Dependent | 2 | 1 | 1 | POSITIVE | POSITIVE | POSITIVE | POSITIVE | POSITIVE | POSITIVE | NEGATIVE | NEGATIVE | NEGATIVE | NEGATIVE |
| | 34b | Non-QE | | 2 | 1 | 1 | POSITIVE | POSITIVE | POSITIVE | POSITIVE | POSITIVE | POSITIVE | NEGATIVE | NEGATIVE | NEGATIVE | NEGATIVE |
| | 34c | Non-QE | Independent | 4 | 1 | NULL | POSITIVE | POSITIVE | POSITIVE | POSITIVE | POSITIVE | POSITIVE | NEGATIVE | NEGATIVE | NEGATIVE | NEGATIVE |
| 35 | 35a | Non-QE | Presumably Dependent | 1 | 1 | NULL | POSITIVE | POSITIVE | POSITIVE | POSITIVE | POSITIVE | POSITIVE | NEGATIVE | NEGATIVE | NEGATIVE | NEGATIVE |
| | 35b | Non-QE | | 1 | 1 | NULL | POSITIVE | POSITIVE | POSITIVE | POSITIVE | POSITIVE | POSITIVE | NEGATIVE | NEGATIVE | NEGATIVE | NEGATIVE |
| | 35c | Non-QE | Independent | 1 | 1 | NULL | POSITIVE | POSITIVE | POSITIVE | POSITIVE | POSITIVE | POSITIVE | NEGATIVE | NEGATIVE | NEGATIVE | NEGATIVE |
